# On the prospect of identifying adaptive loci in recently bottlenecked populations

**DOI:** 10.1101/009456

**Authors:** Yu-Ping Poh, Vera S. Domingues, Hopi E. Hoekstra, Jeffrey D. Jensen

## Abstract

Identifying adaptively important loci in recently bottlenecked populations – be it natural selection acting on a population following the colonization of novel habitats in the wild, or artificial selection during the domestication of a breed – remains a major challenge. Here we report the results of a simulation study examining the performance of available population-genetic tools for identifying genomic regions under selection. To illustrate our findings, we examined the interplay between selection and demography in two species of *Peromyscus* mice, for which we have independent evidence of selection acting on phenotype as well as functional evidence identifying the underlying genotype. With this unusual information, we tested whether population-genetic-based approaches could have been utilized to identify the adaptive locus. Contrary to published claims, we conclude that the use of the background site frequency spectrum as a null model is largely ineffective in bottlenecked populations. Results are quantified both for site frequency spectrum and linkage disequilibrium-based predictions, and are found to hold true across a large parameter space that encompasses many species and populations currently under study. These results suggest that the genomic footprint left by selection on both new and standing variation in strongly bottlenecked populations will be difficult, if not impossible, to find using current approaches.

## Introduction

Identifying the genes driving speciation or adaptation following the colonization of novel habitats is a major focus of both ecological and evolutionary genetics. The rapid fixation of a favorable allele by directional selection results in reduced genetic variability [1] and well-described skews in the frequency spectrum at linked loci via genetic hitchhiking ([2]; and see review [3]). However, demographic factors alone may also produce similar patterns, particularly reductions in population size that subsequently lead to an increased rate of genetic drift. Exploring this issue analytically, Barton [3] demonstrated that a selective sweep had similar effects on neutral diversity as a founder event. In particular, the coalescence events induced by the size reduction, followed by population growth, result in a scenario in which the distribution of neutral genealogies matches that expected under a selective sweep model (for further discussion, see review [4]). Despite this important result, it has nonetheless been proposed that because demographic events affect the entire genome, whereas selective events have only locus-specific effects (*e.g*., [5]), it may be possible to take a simple outlier approach to identify recently selected loci [6]. However, consistent with the analytical results, it subsequently has been demonstrated via simulation that such outlier-based genomic scans based upon neutral equilibrium null models are prone to high false positive rates [4,7,8], owing to an inability to distinguish neutral non-equilibrium models from non-neutral equilibrium models.

To circumvent these difficulties, Nielsen and colleagues [9] proposed the idea of utilizing the background site frequency spectrum (SFS) as a null model in a statistic termed Sweepfinder. In brief, rather than depending upon comparison with the standard neutral model, this class of tests simply would identify putatively adaptive loci that are unusual relative to the background level of genomic variation. With the same notion, but utilizing patterns of linkage-disequilibrium (LD) instead of the SFS as with Sweepfinder, the *ω*_max_ [10] and iHS [11] statistics also have been proposed. Particularly with the emergence of next-generation sequencing, an ever-increasing number of studies have relied on these promising ‘background-effect-based’ approaches – utilizing huge amounts of data to construct the background SFS / LD (thus controlling for demography, in principle) – to identify loci contributing to a local adaptive response (*e.g.,* [12-17]).

Because a great majority of these applications seek to identify adaptively significant loci in severely bottlenecked populations (*e.g*., populations that have recently colonized novel habitats or domesticated species), and in light of Barton’s [18] important analytical results suggesting that the background SFS may not in fact be distinct relative to a swept region in bottlenecked populations, here we revisit the notion that the background SFS may be used to distinguish adaptively important loci in non-equilibrium populations. Thus, building on the results of Pavlidis et al. [10], we directly evaluate the ability of these approaches to (1) identify selected loci within recently bottlenecked populations (rather than considering neutral bottleneck models vs. equilibrium selection models) across a wide-range of bottleneck scenarios, and (2) localize the site of the beneficial fixation.

To test the utility of these approaches, we first focused on two particularly illustrative examples. First, we used the oldfield mouse (*Peromyscus polionotus)* from Florida’s Gulf Coast, in which the selected phenotype (cryptic camouflage; [19]), and its underlying genotype (a single non-synonymous mutation [Arg^65^Cys] in the *melanocortin-1 receptor* [*Mc1r*]; [20]) are well documented. In addition, both the geological age of the islands [21] and the time and severity of the colonization bottleneck have been estimated [22]. Specifically, the derived *Mc1r* allele contributes to lighter camouflaging pigment of the Santa Rosa Island beach mice (*P. p. leucocephalus*) relative to the darkly pigmented, ancestral mainland subspecies (*P. p. subgriseus*) (Figure 1A; [20,23]). Thus, it is reasonable to expect an identifiable selective sweep signal around the *Mc1r* gene using the aforementioned population-genetics approach. However, we were unable to detect any significant signal in *Mc1r* or its surrounding regions by either SFS-based or linkage disequilibrium (LD)-based methods (Figure 1; also see [22]), despite the unusually precise knowledge of recent selection acting on genotype/phenotype.

**Figure 1.**
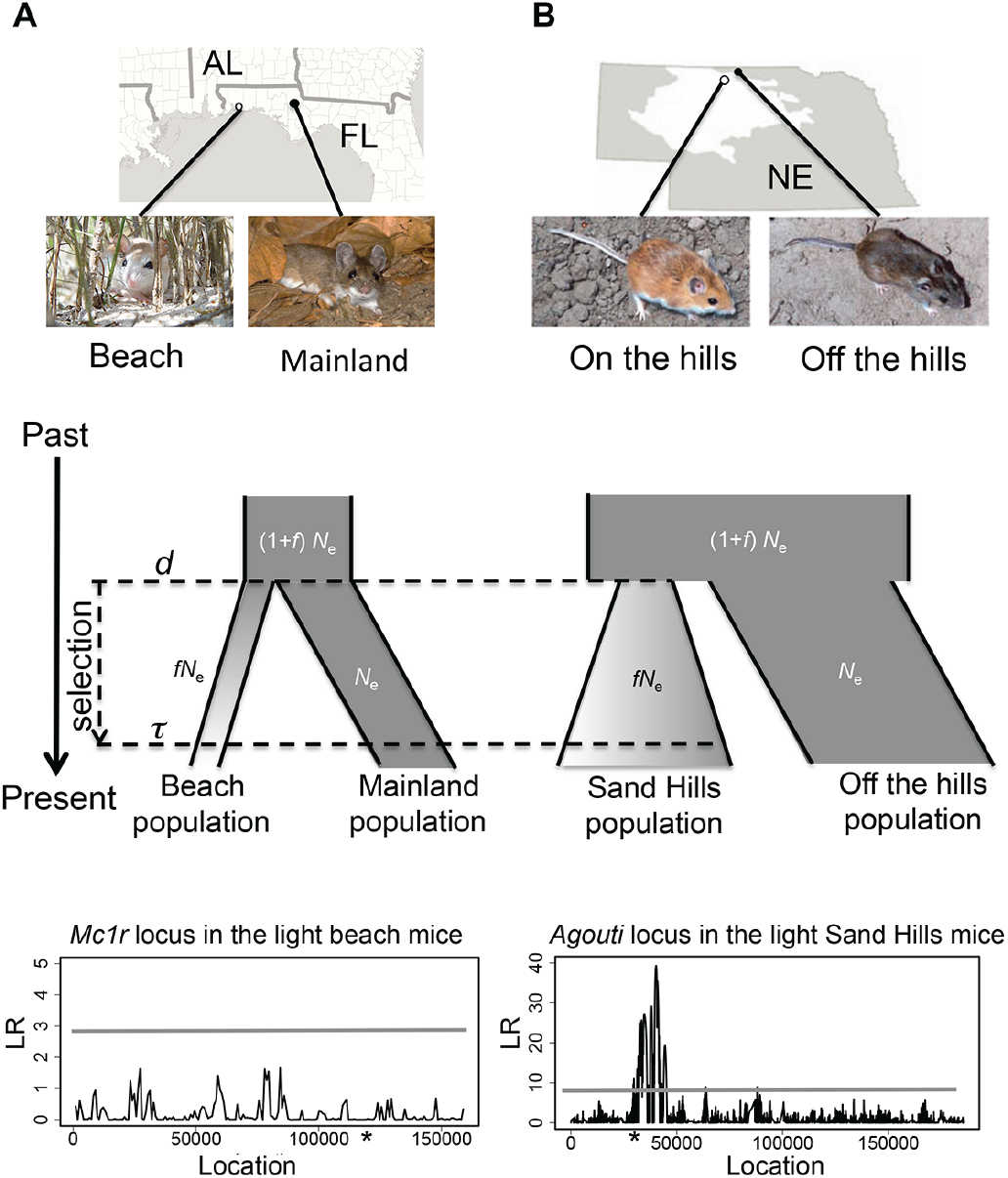
The inferred demographic history of the Florida and Nebraska populations. Geographic location and photos of the derived light and ancestral dark mouse populations from (A) Florida (photos by J. Miller and S. Carey) and (B) Nebraska (photos by C. Linnen) (top panel). Cartoon representation of the inferred demographic model of the two species (middle panel). Both models include selection acting on the bottlenecked population (with effective population size reduced to *fN*_e_, where *N*_e_ is the ancestral population size) immediately after the divergence from the ancestral population at time *d*, and the selected allele becomes fixed at time *τ*. Likelihood ratio (LR) profile of Sweepfinder in both populations of light-colored mice (bottom panel), where the horizontal line indicates the significance cutoff. Stars indicate the approximate location of causal mutations conferring light pigmentation. Because there are multiple *Agouti* alleles, we here polarize (into “light” or “dark” class) based on the SNP mostly strongly associated with pigment variation (as described in [26]).

In a second example, populations of *P. maniculatus* in Nebraska have recently evolved cryptic coloration in a novel light substrate habitat as a result of the formation of the Sand Hills approximately 10,000 years ago [24,25]. The Nebraska Sand Hills mice have accumulated multiple adaptive mutations within the pigmentation locus *Agouti* (Figure 1B). But, unlike in beach mice, Sweepfinder detected large and strong selective footprints around SNPs associated with different pigmentation traits (Figure 1; [26]). This Sand Hills population has experienced a recent population reduction similar in both timing and severity to beach mice. For reference, both these bottlenecks are more extreme than that of human populations out of Africa [27,28]) but comparable to the population reduction associated with dog breed formation [29].

Here, we explore the major factors contributing to this difference in performance between the Florida and Nebraska mouse populations – and more broadly explore the parameter space over which population-genetic approaches may be expected to be successful via simulation. While this study is motivated by the results observed in *Peromyscus* (as this is in many ways a ‘best-case scenario,’ in which selective pressure, phenotype, and underlying genotype are all well described), our results are broadly applicable across systems as the field continues to maintain a strong focus upon identifying locally adaptive loci in strongly bottlenecked populations that are associated with recent colonization (*e.g.,* [30,31], domestication (*e.g*., [32-34], and infection (*e.g*., [35-38]).

## Materials and Methods

### Empirical data analysis

To evaluate the performance of commonly used statistics to detect selective sweeps, we used two well-studied populations of *Peromyscus* mice—one in which signatures of selection were absent and a second in which they were strong—as a starting point. We first utilized the Santa Rosa Island population of beach mice (*P. polionotus*) in which a *Mc1r* variant contributing to cryptic coloration has been fixed [22]. Nineteen individuals from Santa Rosa Island were sampled. The SureSelect capture array (Agilent Technologies, Santa Clara, CA) based on a *Peromyscus Mc1r*-containing BAC clone was designed to enrich the templates for the *Mc1r* locus, and then the capture library was sequenced on an Illumina HiSeq 2000 (Illumina Inc., San Diego, CA) (see [22]). Raw sequence data are available at the NCBI Sequence Read Archive (accession number: SRA050092.2). We used the Burroughs-Wheeler Alignment (BWA) tool to perform mapping and alignment, and used GATK software to call the SNPs and identify genotypes.

The ancestral mainland population size was estimated to be *N*_a_ ∼2500, representing a 99.9% population size reduction associated with the colonization of Florida’s Gulf Coast approximately 3,000 years ago (∼ 7000 generations ago). Then, we applied Sweepfinder on the *Mc1r* genomic region and determined the significance level by *ms* simulation [39] based on the above estimated demographic parameters.

Similarly, 91 individuals of the Nebraska Sand Hills mice (*P. maniculatus*) were collected, and ∼180kb encompassing the *Agouti* locus was sequenced. The sequence data were deposited in NCBI Sequence Read Archive (accession number SRP017939). The SureSelect capture array based on a *Peromyscus Agouti*-containing BAC clone was designed to enrich the templates for the *Agouti* locus. The sequencing and mapping strategies were identical to those used above for *P. polionotus* and *Mc1r*, and further details can be found in [26].

The Sand Hills mice likely colonized the novel light dunes approximately 3,000 years ago at which time they also experienced a severe bottleneck (∼99.6% reduction in population size; [25,26]). Thus, the timing (denoted as *d*) and severity (denoted as *f*) of the bottleneck are remarkably similar in both populations. However, the size of the Nebraska population (*N*_*e*_ = ∼50,000; Table 1) is estimated to be 200 times greater than the Florida population (*N*_*e*_ = ∼2500; Table 1). However, because the derived light phenotype in the Nebraska population is not fixed in the sampled population [see 26] and Sweepfinder and *ω*_max_ are only applicable for complete sweeps, we divided the entire dataset into “light” and “dark” alleles at the *Agouti* locus based on the SNP most strongly associated with pigmentation, in this case, the tail stripe phenotype. Thus, the light alleles represent a population in which the selected allele has been recently fixed, while the dark alleles are used as a reference population characterized by a shared demographic history.

**Table 1.** Comparison of demographic parameters in Florida mice and Nebraska mice

|  | <u>Florida mice</u> | <u>Nebraska mice</u> |
| --- | --- | --- |
| $N_e$ | 2482 | 53080 |
| Estimated $\Theta^a$ | $3.672 \times 10^{-4}$ | $7.792 \times 10^{-3}$ |
| Severity of recent bottleneck<sup>b</sup> | 0.001 | 0.004 |
| Population size recovery<sup>c</sup> | 0.413 | 0.662 |
| Time of bottleneck (in $2N$ generations) | 1.225 | 0.067 |
| Time of bottleneck (in years) | 3040 | 2866 |

<sup>a</sup>pairwise nucleotide diversity per site
<sup>b</sup>ratio of bottleneck population size to ancient population size
<sup>c</sup>ratio of contemporary to ancient population size

### Simulated data analysis

To parameterize both demographic and selection estimates, we performed coalescent simulations using the code of Thornton and Jensen [40], and the parameters used here follow their definitions. In addition to the Florida and Nebraska population models, we performed general simulations to better describe the performance of these statistics. Both comparable (severity of bottleneck, *f* = 0.01) and less severe bottlenecks (*f* = 0.1 and *f* = 0.5) were evaluated in three different population sizes (*N*_*e*_ = 10^4^, 10^5^ and 10^6^). For the simulations, we used the mutation and recombination rates estimated from *Mus domesticus* (*μ* = 3.7 × 10^−8^, [41]; *r* = 5.6 × 10^−7^, [42]). The sample size (*n*; number of chromosomes) = 40 (see Figure S1 for comparisons of different sample sizes) and region length (*L*) = 180kb (see Figure S2 for a comparison of different region lengths) are fixed in all simulated datasets to match the data obtained in the *Peromyscus* populations, thus representing realistic empirically-based parameters.

In our demographic simulations, we considered a selective sweep on a single *de novo* mutation at position 90kb (*i.e*., the middle of the simulated region) with selection coefficients of *s* = 0.001, *s* = 0.01 or *s* = 0.1 arising immediately after the divergence from the ancestral population. These models result in fixation times (*τ*) ranging from 0.01 to 0.3 2*N* generations in the past. The population size reduction occurs immediately after divergence from the ancestral population, and recovers 0.01 2*N* generations prior to sampling. Finally, 100 replicates for each model were generated and analyzed using the commonly used background SFS approach (Sweepfinder; [9]) as well as the sliding window LD method (*ω*_max_; [10]). Significance cutoffs were determined via neutral simulation in *ms* [39], with the demographic model and *θ* fit to each case. Following Nielsen and colleagues [8], the 95th percentile of the statistic *Λ*_*SF*_ denotes the threshold value. Given that the expected size of the sweep region can be approximated as 0.01 *s*/*r* base pairs [43], the footprints of selection should be captured within 10kb window surrounding the selected site (see Figure S3 for the empirically observed LD decay). We thus considered the rejections of neutrality within 10kb as true positives (TP), and those outside the targeted region as false positives (FP). The TP and FP rates were used as the major indicator for the performance of Sweepfinder and *ω*_max_ to identify selective sweeps.

### Results & Discussion

The likelihood profiles of Sweepfinder in both the Florida beach mice and Nebraska Sand Hills mice are given in Figure 1, highlighting a significant result only in the Nebraska population. To investigate this finding, we first performed a series of simulations using demographic models mimicking the population history of the Florida and Nebraska mice (Table 1), accompanied by a single hard sweep. Here we assume that selection began at the time of the split from the ancestral population, and the selected allele was fixed at 0.1 or 0.3 2*N* generations ago, with strengths ranging from 0.001 to 0.1. The observed median values of polymorphism in the replicates range from *π =* 6.5 × 10^−5^ – 6.9 × 10^−5^ for Florida-model mice and *π =* 6.6 × 10^−5^ – 1.1 × 10^−4^ for Nebraska-model mice, with the SFS skewed towards rare alleles (with observed median values of Tajima’s *D* = -1.80 – -2.13).

For small population sizes (*i.e.,* the Florida population), we found that Sweepfinder has very limited power to detect recent selective fixations (Figure 2), while TP improves for larger population sizes (*i.e.,* the Nebraska population) – though this increase in power is also associated with an increased FP rate (Table 2). Similarly, the *ω*_max_ statistic is not able to clearly discriminate the selected loci from the neutral background when the population size is small, but again TP improves as population size increases. As has been previously described (*e.g.,* [44]), power diminishes quickly as the time since fixation (*τ* given in 4*N* generations) increases – with Sweepfinder failing to detect any rejections of neutrality for *τ*= 0.3. At *τ* = 0.1, the power of Sweepfinder and *ω*_max_ are comparable. In general, the rejection rate (TP and FP) of Sweepfinder is lower than *ω*_max_ in both examples, though higher FP in many ways presents a greater concern. Thus, the successful empirical identification of the signature of selection in the Nebraska Sand Hills mice, relative to the Gulf Coast population, by Sweepfinder likely is attributable to the larger population size.

**Figure 2.**
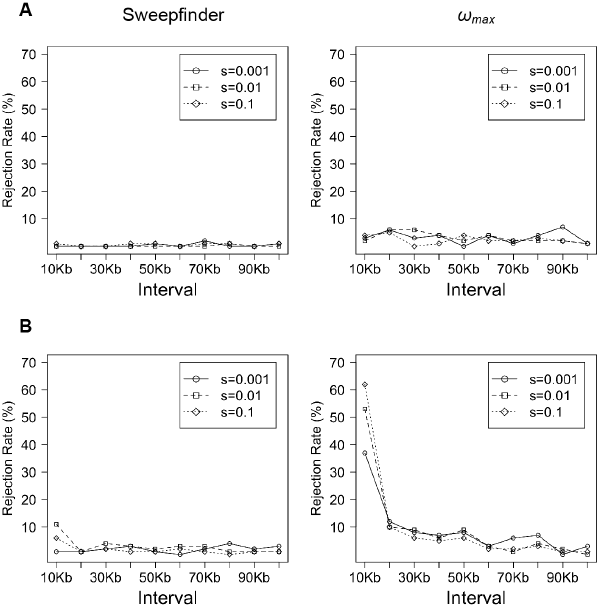
The fraction of simulated replicates rejecting the neutral model by Sweepfinder and *ω*_max_, with varying population size. The simulations with demographic models mimic the history of (A) Florida beach mice (*N*_e_ = 2500) and (B) Nebraska Sand Hills mice (*N*_e_ = 50,000). The time of the bottleneck (*t*_r_ = 0.1) and time since fixation *τ* = 0.1) are fixed, but selection strength varies from *s* = 0.001 to 0.1. Ideal performance would be indicated by all replicates showing a significant signal at very small window sizes, suggesting an ability to localize the target.

**Table 2.**
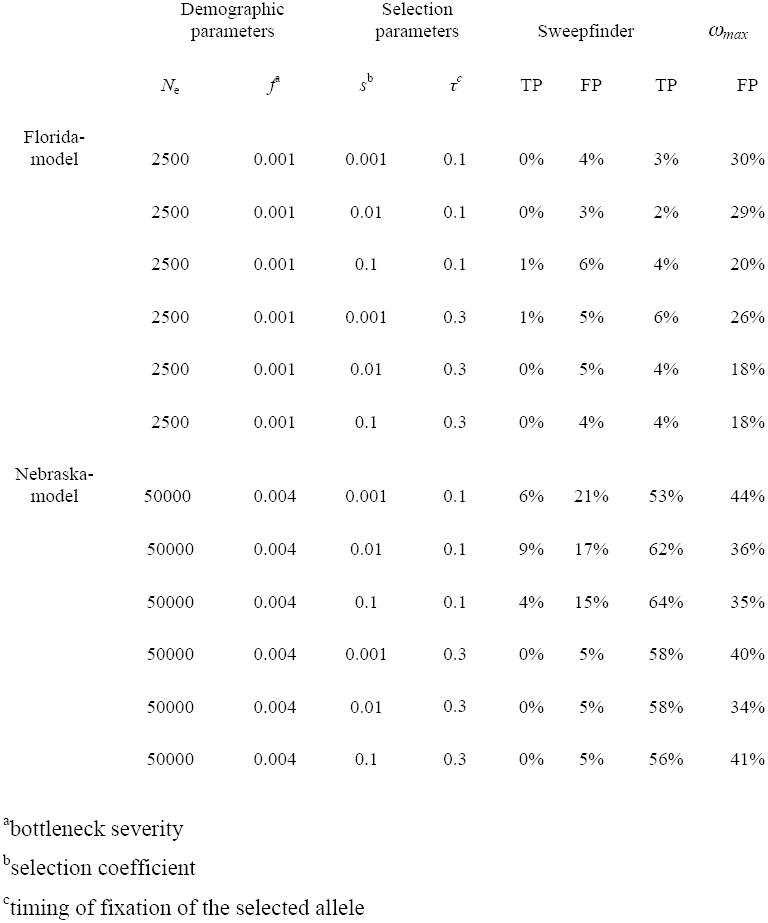
The true positive (TP) and false positive (FP) rates of Sweepfinder and *ω*_*max*_ in Florida and Nebraska population models. A TP is defined as a significant rejection within 10kb of the true target.

To consider more generalized parameters, we examined performance across simulations of varying *N*_e_ and *f* (Figure 3). In general, the TP rate of Sweepfinder is higher than *ω*_max_ for populations of small *N*_*e*_ (though both approaches perform poorly), but *ω*_max_ performs better when *N*_*e*_ > 10^5^ (i.e., for *N*_*e*_ = 10^5^, TP ∼ 50%; for *N*_e_ = 10^6^, TP ∼ 60%), despite the severity of the bottleneck. The improved performance of *ω*_max_ is related to the increasing SNP density, which increases for larger *N*_e_. However, the TP rate of Sweepfinder remains relatively constant across varying *N*_e_ or *f* (TP ∼ 10%). To further explore the effect of the timing of selection, we compared the Sweepfinder and *ω*_max_ results for a range of times since fixation (Figure 4). As suggested by the mouse examples above, Sweepfinder has no power to reject neutrality when a beneficial fixation is older than 0.01 2*N* generations. Similarly for *ω*_max_, power is maximized when the sweeps are recent and occur in large populations.

**Figure 3.**
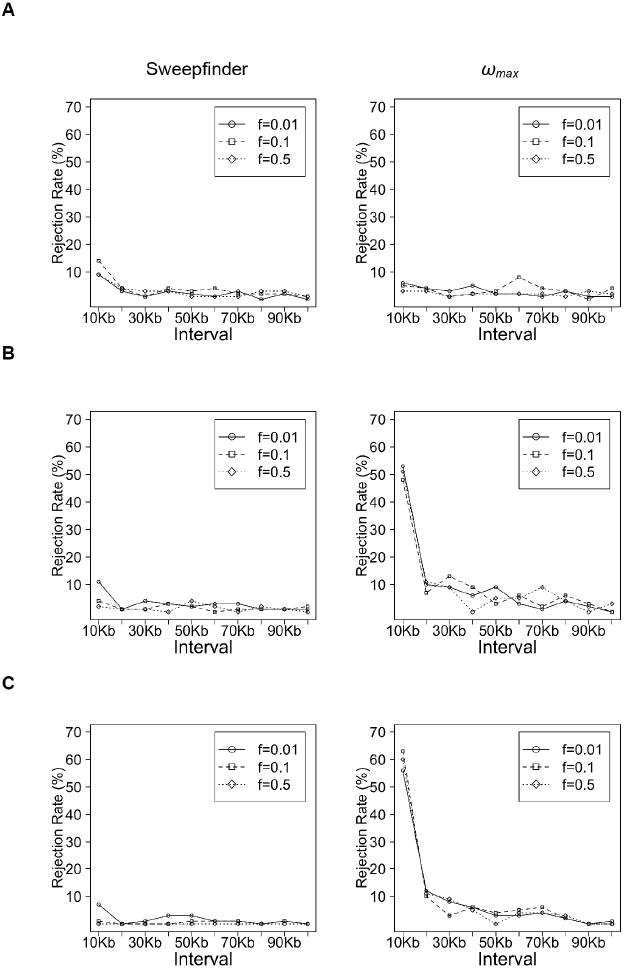
The fraction of simulated replicates rejecting neutrality by Sweepfinder and *ω*_*max*_, with varying bottleneck severity. Simulations with ancestral population size equal to (A) 10^4^, (B) 10^5^ and (C) 10^6^. Selection strength (*s* = 0.01), time since fixation (*τ* = 0.1), and time since bottleneck (*t*_r_ = 0.1) are fixed, but bottleneck severity (*f*) varied from 0.01 to 0.5.

**Figure 4.**
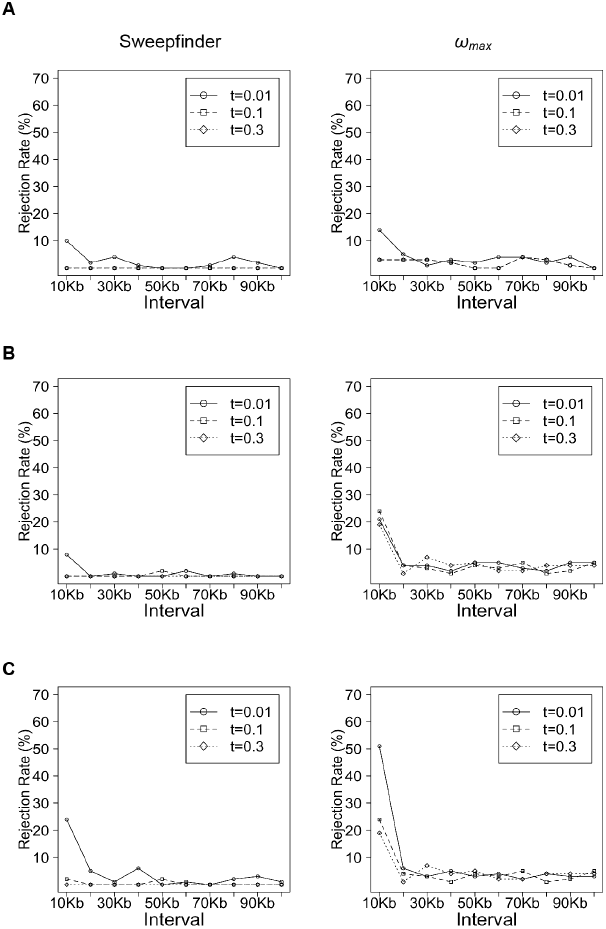
The fraction of simulated replicates rejecting neutrality by Sweepfinder and *ω*_*max*_, varying the time since the beneficial fixation. Simulations with ancestral population size equal to (A) 10^4^, (B) 10^5^ and (C) 10^6^. Selection strength (*s* = 0.01) and time since bottleneck (*t*_r_ = 0.01) are fixed, but the time of selected allele fixation (*τ*) varied from 0.01 to 0.3.

The results from both the empirical examples and the more general simulations together highlight two fundamental lessons. First, the skew in the SFS associated with a selected region is not unusual relative to the background genomic patterns under a variety of bottleneck models owing to the fact that the coalescent processes underlying both the selected locus as well as the surrounding neutral loci are similar, as described by Barton [45]. Second, LD-based expectations generally outperform SFS-based expectations under these models (particularly for large population sizes), supporting the theoretical predictions of Stephan *et al*. [46] in describing the advantages of this specific post-fixation LD expectation (*i.e.,* elevated LD flanking the beneficial mutation, but reduced LD spanning the site), further highlighting the value of generating linkage information, rather than simply SNP frequencies, in future genomic studies [47]. Importantly however, even this LD pattern is not exclusive to selective sweeps, and also may be generated under certain neutral bottleneck models.

## Conclusions

The ability to detect the footprint of a selective sweep in genomic data from bottlenecked populations remains as an important and largely unresolved challenge. The results presented here strongly suggest that the widely utilized approach of employing the background SFS as a null model has not much improved our ability to identify true selective sweeps for much of the parameter space of interest to biologists. Troublingly, the false positive rate found by these models is often in excess of power, suggesting that the majority of significant results in such populations are likely erroneous. In the extreme case of beach mice – in which the target of selection has been functionally validated – we have not identified any existing population-genetic-based statistic capable of identifying this causal variant. In comparison, the successful identification of beneficial mutations in Nebraska mice can be attributed to its larger population size as well as the recurrent and recent selective events still ongoing in the Sand Hills population. Thus, these data underscore both a need for great caution when interpreting results from selection studies in recently bottlenecked populations and for continued methodological and theoretical development, specifically inference procedures capable of jointly estimating selection and demography simultaneously.

---

**Figure S1.**
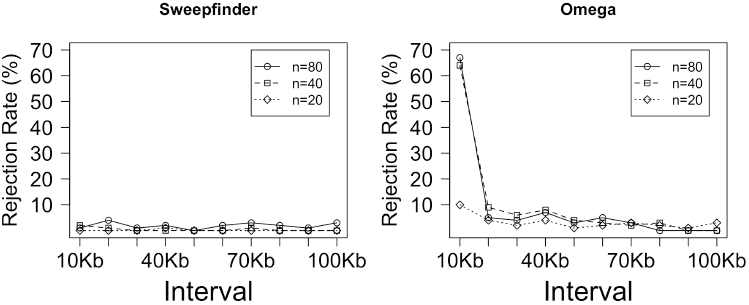
The fraction of simulated replicates rejecting the neutral model by Sweepfinder, with varying sample size. The simulations with demographic models mimic the estimated history of Nebraska Sand Hills mice (*N*_e_ = 50,000), the time of the bottleneck (*t*_r_ = 0.1), the time since fixation (*t* = 0.1), and the selection strength (*s* = 0.1) are fixed. Here, the sample size varies from n = 20 to 80. Ideal performance would be indicated by all replicates showing a significant signal at very small window sizes, suggesting an ability to localize the target.

**Figure S2.**
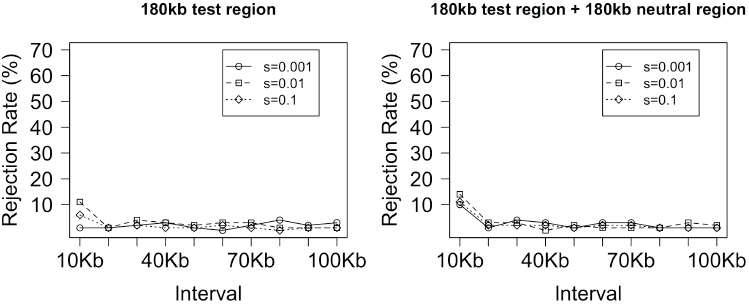
The fraction of simulated replicates rejecting the neutral model by Sweepfinder, with and without another 180kb simulated neutral region. The simulations 180kb with demographic models mimic the estimated history of Nebraska Sand Hills mice (*N*_e_ = 50,000), the time of the bottleneck (*t*_r_ = 0.1), the time since fixation (*t* = 0.1), but selection strength varies from *s* = 0.001 to 0.1. The right panel shows the Sweepfinder performance with another 180kb simulated neutral regions with the same demographic parameters added. The results suggested that Sweepfinder could gain more efficacy in identifying sweeps with more neutral SNPs to build the background SFS, but the improvement is modest.

**Figure S3.**
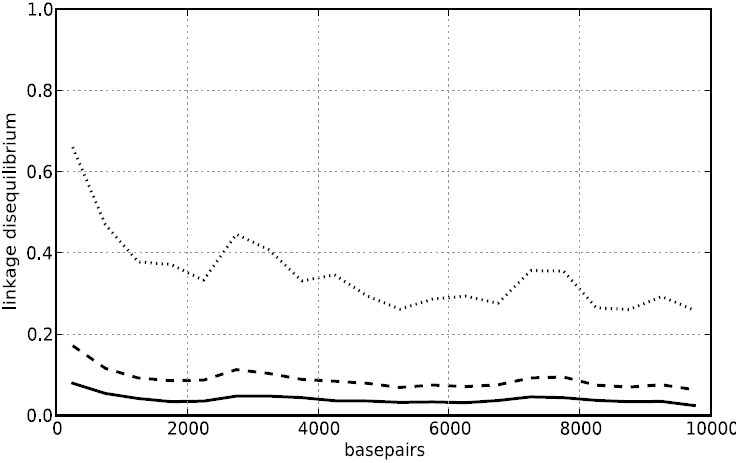
Decay of linkage disequilibrium (LD) as a function of physical distance between variable sites. In all panels the solid lines represent medians for each X axis category (physical spacing bin) centered on the plotted X coordinate, dashed lines represent means of spacing bins, and dotted lines represent 95th percentiles of spacing bins.

